# Same-sex sexual behaviour reduces reproductive success but mitigates aggression in a subsocial insect

**DOI:** 10.64898/2026.10.18.759864

**Authors:** Nicholas E. Parlamis, Jakob Wiil, Natalie Pilakouta, Nathan W. Bailey

## Abstract

Same-sex sexual behaviour (SSB) is widely observed across taxa, yet its evolutionary maintenance remains debated and its fitness consequences remain surprisingly understudied. Here, we quantified the fitness consequences of male SSB in the burying beetle *Nicrophorus vespilloides* and tested three prominent hypotheses about its function: (*i*) that it provides practice for future mating, (*ii*) that it reduces tension through dominance assertion, and (*iii*) that it arises indirectly from indiscriminate mating. Males were observed in same-sex pairings to quantify SSB and then paired with females to measure their subsequent mating behaviour and reproductive success. Same-sex sexual behaviour was common, occurring in 55% of same-sex pairs, and was associated with reduced reproductive success driven by higher brood failure rates. Both males that initiated, and those that received, mounting attempts from other males suffered reduced fitness. Contrary to the practice hypothesis, SSB did not increase mating speed with females and was associated with reduced mating frequency. However, SSB was associated with reduced aggression, supporting the hypothesis that SSB reduces tension through dominance establishment. Our findings show that SSB can reduce reproductive success but may be maintained due to a beneficial reduction in aggression.

## 1. Introduction

Same-sex sexual behaviour (SSB) involves sexual, reproductive, or, under some definitions, parental behaviours between two individuals of the same sex (Bailey & Zuk, 2009; Bagemihl, 1999; Sommer & Vasey, 2006). Evolutionary explanations for this taxonomically diverse trait are numerous but can mainly be split into adaptive and nonadaptive hypotheses (Bailey & Zuk, 2009). Adaptive hypotheses suggest that SSB provides a net direct fitness benefit that allows the behaviour to persist via positive selection. These include social cohesion (Mann, 2006; Clive *et al.,* 2023), kin selection (Rahman & Hull, 2005), overdominance (Gavrilets & Rice, 2006), sexually antagonistic selection (Camperio-Ciani *et al*., 2004), practice for future mating (McRobert & Tompkins, 1988), tension-reduction in aggressive encounters (Green *et al*., 2026), and social dominance assertion (Vervaecke & Roden, 2006; Lane *et al*., 2016). Non-adaptive hypotheses propose that SSB is costly to the actor but persists due to indirect associations with other traits; in this view it is sometimes framed as a byproduct of selection or a result of balancing selection. Such hypotheses include mistaken identity arising from incomplete sex discrimination ability (Switzer *et al*., 2004; Bailey & French, 2012; Lerch & Servedio, 2020), the so-called ‘prison effect’ (Van Gossam *et al*., 2005; Bonnet *et al*., 2016), and overdominance or sexually antagonistic selection (Gavrilets & Rice, 2006; Hoskins *et al.,* 2015).

Under both scenarios, research on SSB is often predicated upon the assumption that it incurs fitness costs due to missed mating opportunities (e.g., Gavrilets & Rice, 2006; Levan, 2009). This assumption has recently been challenged in vertebrates, where there is evidence that SSB decreases different-sex sexual behaviour (DSB) to a negligible degree and has no negative effects on an individual’s reproductive success (e.g., *Macaca mulatta*; Clive *et al.,* 2023). Similarly, same-sex female pair bonds and sexual interactions in some birds, such as the Laysan albatross (*Phoebastria immutablis*), increase fitness in circumstances where male mates are scarce (Young *et al*., 2008; Young & VanderWerf, 2014). There is arguably less data about the reproductive fitness consequences of SSB in invertebrates. However, a study by Berger *et al*. (2016) found that male SSB in seed beetles (*Callosobruchus maculatus*) is genetically associated with increased female, but decreased male, reproductive success, with the pattern reversed for female SSB. Given the lack of information about the fitness effects of SSB in invertebrates, the assumed ‘inherent’ costs of SSB are mainly extrapolated from known costs of DSB. These are either direct energetic costs, such as the energy invested in reproductive behaviours (Scharf *et al.,* 2013), or trade-off costs, such as reduction in immune performance (Lawniczak *et al.,* 2007), exposure to predators and parasitoids, which exploit sexual traits such as chemical signals and bioacoustics (see examples in Kotiaho, 2001), and risk of injury (e.g., antennae breakage; Benesh *et al.,* 2007; wing/genitalia damage; Kuriwada & Kasuya, 2009). However, some of this DSB-associated risk of injury arises due to aggressive encounters with rival-males, which SSB has been suggested to de-escalate (e.g., Green *et al.,* 2025).

In this study, we test the reproductive fitness consequences of male SSB in the burying beetle *Nicrophorus vespilloides*, which has been shown to exhibit high levels of SSB (Figure 1A; Engel *et al*, 2015; Morelle, *et al.,* 2026). We measured the reproductive fitness costs or benefits of previously-expressed SSB using sequential, dyadic encounters, first between same-sex individuals and then between different-sex individuals. Analysing all individuals’ SSB as well as aggressive behaviours in these encounters enabled us to evaluate support for several hypotheses about its function (Figure 1B). Most studies on invertebrates invoke non-adaptive hypotheses centered on mistaken identity mechanisms, or as has been more recently proposed, broad mating filters, to explain their findings (Scharf & Martin, 2013; Richardson & Zuk, 2023). However, this is increasingly challenged by research demonstrating that SSB in insects can occur despite accurate sex recognition (Mizumoto *et al*., 2022; Green *et al*., 2025; Bailey & Green, 2026; Ouadah *et al*., 2026).

**Figure 1.**
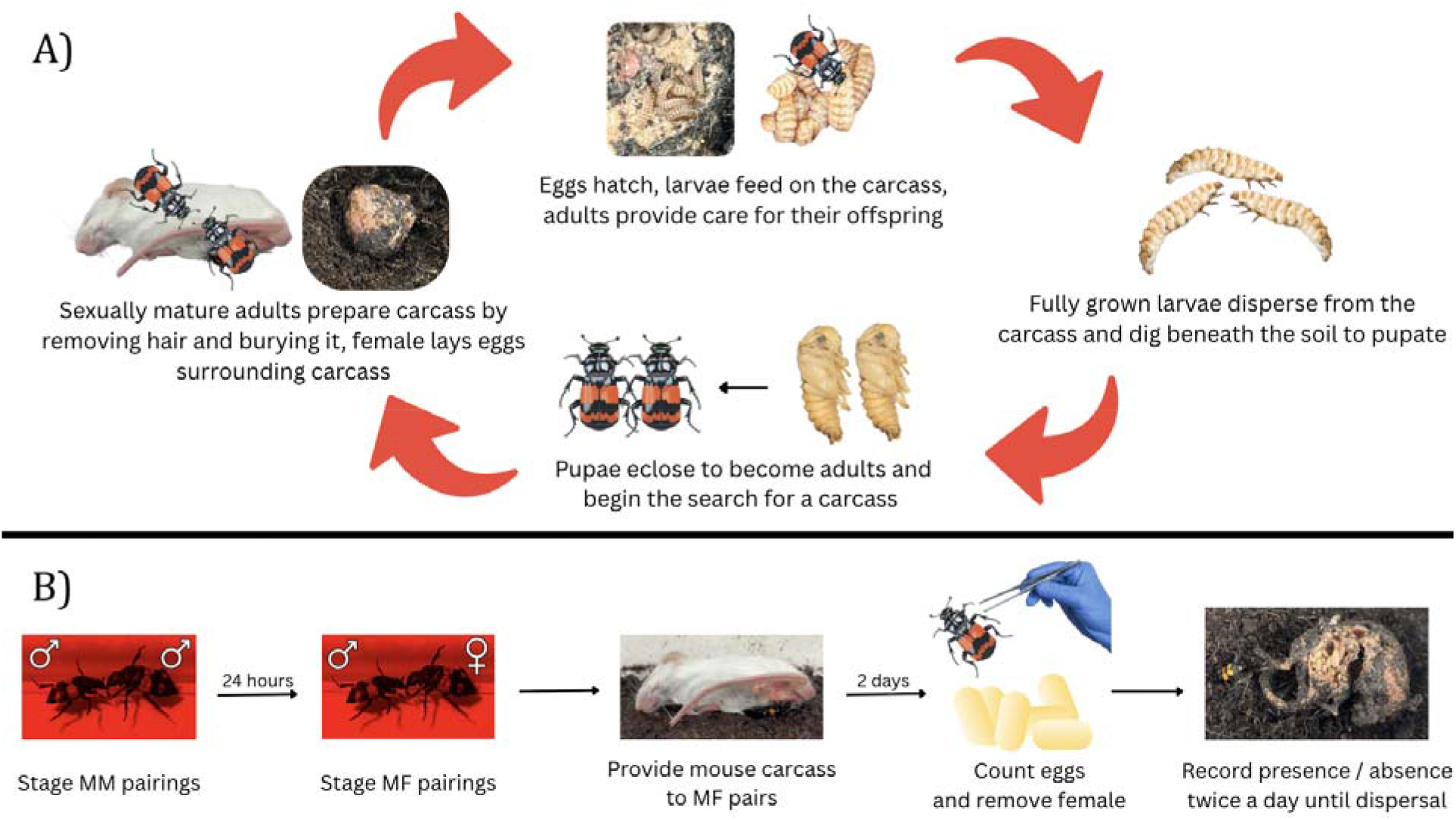
Overview of *Nicrophorus vespilloides* as a research model for studying same-sex sexual behaviour. (A) Life cycle as described by Scott (1998). (B) Illustration of experimental design; see Main Text for details.

*Nicrophorus vespilloides* has previously been used to test whether flexible mate acceptance thresholds can explain SSB. For example, a study by Engel *et al*. (2015) suggested that imperfect sex discrimination favours flexible mate acceptance thresholds for male-male matings, depending on the costs of accidentally rejecting an opposite-sex mating opportunity. Inherent in the study design of Engel *et al*. (2015) and subsequent literature that reframes this mechanism as an adjustable mating filter (Richardson & Zuk, 2023) is that fitness costs for male SSB are low relative to the fitness cost of foregoing a viable mating. However, the direct fitness costs of SSB have yet to be tested and are likely to involve a complex dependency on this species’ particular life history. First, *N. vespilloides* require a carcass for reproduction (Figure 1A), but they will still mate in the absence of a carcass because females can store fertile sperm in their spermathecae (Müller & Eggert, 1989; Scott, 1998). Second, males can mat repeatedly, suggesting they do not become severely sperm limited (House *et al.,* 2008). These two features of burying beetle behaviour may explain the appeal of SSB explanations that rely on loose sex discrimination. However, a lack of evidence for adaptive explanations does not equate to evidence of a lack of adaptive value for SSB. There is evidence, for example, that *N. vespilloides* have sexually dimorphic chemical compositions of cuticular compounds (Steiger *et al.,* 2007), and that they use these to distinguish between the sexes (Keppner *et al.,* 2016). Additionally, males release a sex-specific pheromone to attract partners, even if no carcass is nearby (Müller & Eggert, 1987). Finally, males have fine-tuned chemical recognition abilities and are able to discriminate between virgin and non-virgin females during mating (Steiger *et al*., 2008). Combined with the lack of information about the potential fitness consequences of SSB, these factors call into question the biological plausibility of explanations that rely on mate acceptance errors and argue for detailed investigation of adaptive hypotheses for SSB in this species (Table 1).

**Table 1.** Hypotheses and predictions about the evolved function of male SSB in *Nicrophorus vespilloides* that we tested in this study.

| Hypotheses | Predictions | References |
| --- | --- | --- |
| **Adaptive** |  |  |
| SSB increases fitness by providing practice for future mating encounters | *Individuals initiating SSB will show improved mating success in later encounters with females, via faster initiation of mounting or more mounting events* | Zawistowski & Richmond, 1985; McRobert & Tompkins, 1988; Mann, 2006 |
| SSB increases fitness by reducing costly aggression | *Individuals initiating or receiving SSB will show less aggression than non-SSB males* | Fruth & Hobmann, 2006; Green *et al.*, 2026; Lane *et al.*, 2016 |
| **Non-adaptive** |  |  |
| SSB is costly and arises from a lack of sex discrimination | *Males will attempt to mate females as commonly as with males, and the occurrence of sexual behaviours will not differ between SSB and DSB contexts* | Switzer *et al.*, 2004; Bailey & French, 2012; Lerch & Servedio, 2020 |

## 2. Methods

### (a) Animal husbandry

This study was performed on third- and fourth-generation beetles from a large, outbred laboratory population established from wild-caught beetles originally collected at Bonnytown Farm, UK (56°18’20.8“N, 135 2°44’02.4“W) and Tentsmuir National Nature Reserve, UK (56°25’44.7“N, 2°52’11.2“W) in August–September 2025. Beetles were housed individually in transparent plastic containers (12×8×2 cm) filled with moist soil (Clover Grow brand potting soil) and kept in a dark incubator at 20°C (LMS Series 1A Model 280 NP) to simulate underground conditions. All non-breeding adults were fed small chunks of organic beef and their soil was sprayed with water twice a week. Experimental individuals were virgins, and all pairings occurred within two weeks of reaching sexual maturity (10–24 days after eclosion). Sex was determined at eclosion, and all beetles were re-examined prior to pairing to avoid misidentification.

### (b) Phase 1: Male-male pairings

All pairings occurred between two individuals whose age was no more than eight days apart, and that had no common ancestors for at least two generations. When setting up male-male pairs, all beetles that we intended to be paired that day were taken out of the incubator, one haphazardly selected male from each pair was marked with a small dot of Tipp-Ex in the center of their pronotum, and then all pairs were returned to the incubator to await their trial while the Tip-Exx dried. We also recorded the mass of each individual using an Ohaus PR124 analytical balance (precision of ±0.0001 g) and measured their pronotum width using digital calipers.

All experimental pairings took place in a clear plastic box (17 × 12 × 6 cm) filled with approximately 0.5 cm of moist soil and viewed under red light. Experimental pairs were placed into the box at the same time and then observed over the next 30 minutes via focal sampling for 15 second intervals each minute (Figure 1B). This allowed 4 pairs to be observed at once. This method was chosen instead of continuous or scan sampling because the pilot experiment showed that many interactions were short and likely to be missed by scan sampling. A total of 56 male-male pairs were observed (*n* = 112 individual males). The behaviours measured in this study are described in Table 2. More than one behaviour could be observed within a single 15-second interval (e.g. biting and grappling). However, not-interacting, walking, and hiding were only recorded if no other behaviour occurred, as these were seen as “baseline” states. Immediately following every male-male pairing, Tipp-Ex was applied to the pronotum of the unmarked male to ensure that all males being paired with females were marked, standardizing any effect that Tipp-Ex or its application might have on future male-female pairings, as well as aiding in individual recognition during DSB observations.

**Table 2.** Ethogram of *Nicrophorus vespilloides* behaviours scored during dyadic male-male and male-female interactions.

| Behaviour | Description |
| --- | --- |
| Mounting | Focal individual moves on top of an interacting individual with bodies aligned laterally, abdomen pulsing, and extension of the aedeagus |
| Grappling | Focal individual touches, moves on top of, or pushes the interacting individual with bodies unaligned and no extension of the aedeagus |
| Biting | Focal individual's mandibles clasp any part of the interacting individual, often accompanied by sudden jolts or flipping |
| Stridulating | Focal individual stridulates (audible) |
| Walking | Focal individual moves around the box or attempts to climb the walls |
| Not interacting | Focal individual remains stationary on the floor of the box |
| Hiding | Focal individual buried beneath the soil or actively digging in the soil |
| Wing movement or self-grooming | Focal individual extends or retracts wings, attempts to fly, or cleans antennae, elytra, or any other part of the body |
| Antennal touching | Focal individual's antennae contact an interacting individual, often patting the individual back and forth with each antenna |

### (c) Phase 2: Male-female pairings

Eighty-three of the initial 112 males entered the second phase of the experiment (to control experimental feasibility, some non-SSB males were excluded via haphazard selection). Prior to male-female pairings, all male subjects were classified into three groups based on their behaviour in the male-male pairings. All surviving males that had engaged in SSB (*n* = 55), were classified as either SSB-mounter (i.e. they mounted their partner; *n* = 27) or SSB-mounted (i.e. they had been mounted by their partner; *n* = 28). Mounting was reciprocal during one male-male trial (see Results). Therefore, two males were classified as both SSB-mounters and SSB-mounted. Males that did not engage in SSB were labelled non-SSB (*n* = 29). They were returned to their individual containers in the incubator for 24 hours without being fed, then paired with females and observed under red light for another round of 30-min behavioural observations.

### (d) Phase 3: Reproductive success

Following the male-female behavioural observations, male-female pairs were moved into a new plastic box (17 × 12 × 6 cm) filled with approximately 1cm of moist soil and given a freshly thawed mouse carcass (*Mus musculus*; sourced from Livefoods Direct Ltd) of a standardised size (23±3 g). Pairs were left with the carcass for two days. On the morning of the third day, eggs were counted for each pair, and female beetles were removed from the box to isolate male effects. To quantify parental care duration, we recorded the presence or absence of the male daily at 09:00 and 16:00 under standard laboratory lighting conditions (white light). The male was counted as present if observed in the crypt area (i.e. on the carcass or in the area immediately next to the carcass). If the male was absent three times in a row, he was removed from the box to prevent infanticide and weighed (Oldekop *et al.,* 2007). When the larvae reached the dispersal stage, defined as the point where any last-instar larvae have left the crypt, we counted the number of surviving larvae, and measured total brood mass and the post-breeding mass of the male parent.

### (e) Statistical analyses

Data were analysed and visualised in R v4.4.3 using the packages “dplyr”, “emmeans”, “stringr”, “tidyr”, “tidyverse”, “ggeffects”, and “ggplot2” (Wickham, 2016; Lüdecke, 2018; Wickham *et al.,* 2019; Wickham, 2023; Wickham *et al.,* 2024; Lenth, 2025; Wickham *et al.,* 2026). In male-male trials, we were interested to determine the downstream effects of male SSB in those trials where it occurred. The frequencies of all behaviours (Table 1) were assessed before and after mounting in male-male pairs to test whether previous same-sex mounting predicted behavioural changes. To test the likelihood of displaying behaviours before versus after mounting, a binomial generalized linear mixed-effect model (GLMM) with a binomial error distribution and logit link function was fitted using the “glmer” function in the R package “lme4” (Bates *et al*., 2015). This model was run for each individual focal behaviour and tested for a difference in the probability of each focal behaviour occurring between the pre-mounting and post-mounting periods. In this case, the probability of each behaviour occurring was modelled on the log-odds scale as a proportion of the total observed behaviours for an individual in that period. We were also interested in contrasting these behavioural probabilities with behaviour expressed in trials where no male-male mounting occurred. To do this, we ran an identical model for the non-SSB trials centered around an “anchor time” of t = 10 min (the mean of SSB mounting times in trials where it occurred). This allowed for comparison with the SSB group and provided information on whether any changes in probability of expressing behaviour were merely attributable to typical changes in behaviour patterns over the course of the 30 minute trial, or whether mounting itself was central to the effect.

A Kaplan-Meier survival time analysis was used to examine differences in the speed at which males mounted females depending on SSB status and role, as the data were heavily right-censored. The right-censoring arises because observations of no mounting within 30 minutes do not mean that the individuals would never display mounting if given infinite time. For these analyses, the packages “survival” and “survminer” were used (Therneau, 2024; Kassambara *et al.,* 2026). Reproductive success was analysed by first generating a z-composite measure which accounted for mean larval mass, brood size, and brood survival. To justify the creation of the z-composite, we used the packages “lavaan” and “psych” in a factor analysis and examined the loadings of each variable on the composite (Tables S2 & S3; Rosseel, 2012; Revelle, 2026). Z-composite scores were then compared using Welch’s t-tests between groups. A Fine-Gray competing risks cumulative incidence function model was used to examine brood dispersal times, with brood death as a competing event using the package “cmprsk” (Gray, 2024).

To test whether aggressive behaviour differed by pairing type (male-male vs. male-female) and by engagement in SSB (SSB vs. non-SSB), we fit a generalized linear mixed model (GLMM) with a Poisson error distribution and log link, using the “glmer” function in the “lme4” package in R (Bates *et al*., 2015). The model included pairing type, SSB, and their interaction as fixed effects, as well as individual ID as a random effect to account for repeated observations per individual. Model parameters were estimated via maximum likelihood using the Laplace approximation.

## 3. Results

### (a) Summary of behaviour and offspring production

We observed SSB in 31 out of 56 male-male pairings (55%) during the 30-minute observation period, with 10 pairs displaying more than one instance of mounting (mean instances ± SE: 2.25 ± 0.51). Mounting was always unidirectional (one individual exclusively displaying mounting and the other exclusively being mounted), except for one pair where reciprocal mounting was observed. We observed DSB in 59 out of 83 male-female pairings (71%), with 55 pairs displaying more than one instance of mounting (mean ± SE: 3.47 ± 0.20). The sex of partners did not impact the probability of mounting occurring: there was no statistically significant difference between the proportion of pairings with mounting in male-male pairs versus male-female pairs (55.4% vs 71.1%; χ² test, χ²(1) = 3.62, *p* = 0.057). Additionally, for individuals who displayed both SSB and DSB mounting (*n* = 20), whether the mounted individual was male or female made no difference to the duration of mounting (Welch’s t_48.17_ = −1.87, p = 0.067)

We found variation in patterns of aggressive behaviour depending on whether males experienced SSB or not during their male-male trials (Figure 2). Aggression accompanied SSB in male-male trials. Male-male pairs in which SSB was expressed by either partner showed significantly higher levels of aggression (mean count of grappling or biting = 2.52 instances ± 0.28) compared to non-SSB pairs (mean count = 0.73 instances ± 0.14; Wilcoxon rank-sum test, W = 673, *p* < 0.001). In male-female pairs, male aggression towards females did not depend on males’ previously experienced SSB (Wilcoxon rank-sum test, W = 789 *p* = 0.74). However, this is due to opposing trends for SSB and non-SSB males. For non-SSB males, aggressive behaviour counts were 294% higher in male-female pairings than in male-male pairings (post-hoc pairwise EMM contrast, rate ratio = 3.94, *p* < 0.0001). In contrast, for SSB males, the rate of aggression in male-female pairings was 26% lower than in male-male pairings (rate ratio = 0.74, *p* = 0.024). Thus, the prior experience of SSB was associated with reduced aggression towards females in subsequent mating encounters (Figure 2). Put another way, SSB individuals were less aggressive when paired with females than they were when previously paired with males. However, non-SSB individuals showed the opposite pattern, with higher aggression towards females they were paired with than they showed in previous encounters with other males. This resulted in similar aggression levels displayed by SSB and non-SSB individuals in female-female pairs.

**Figure 2.**
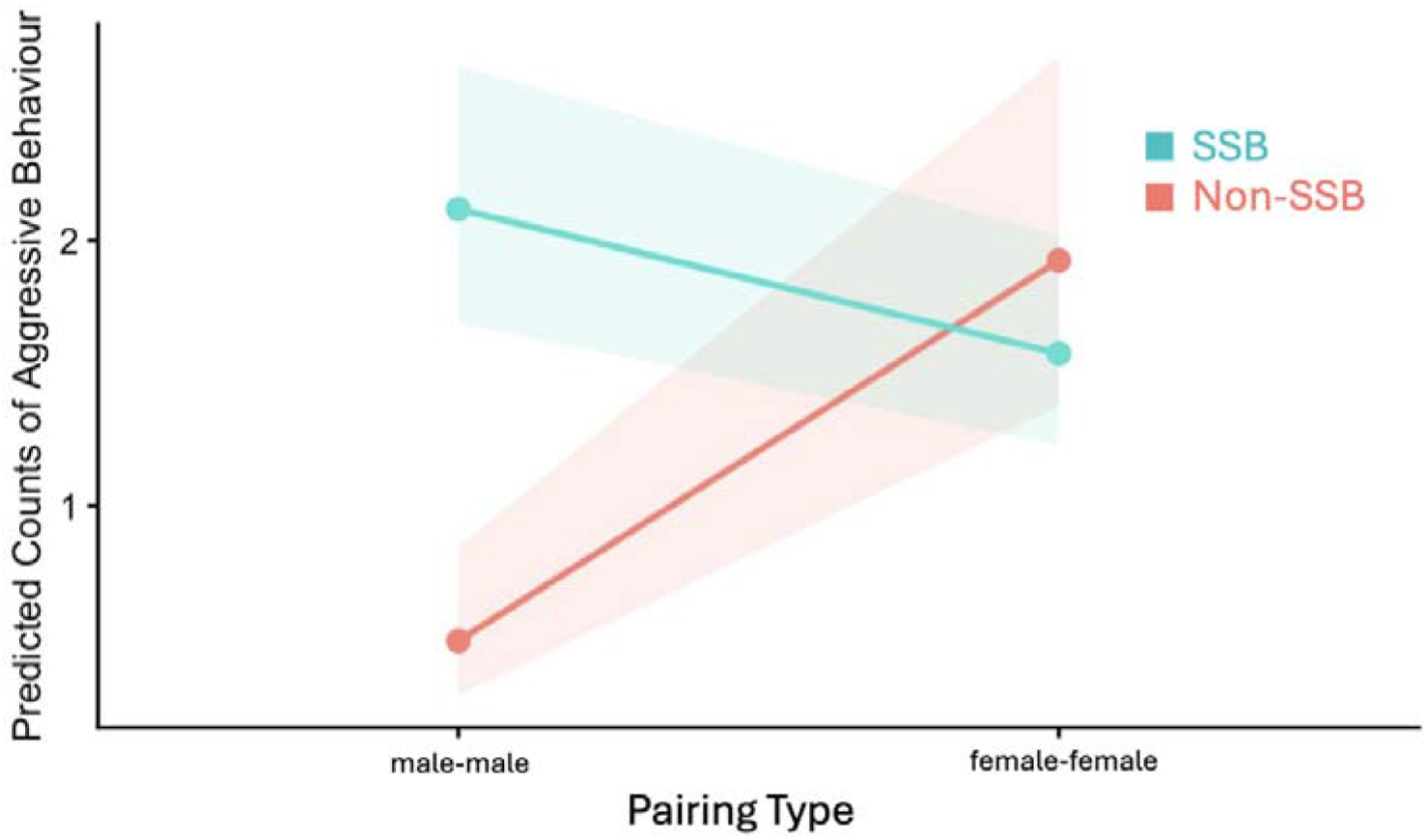
Effect of SSB expressed during male-male encounters on male-male aggression and on intersexual aggression during subsequent male-female encounters. The interaction between SSB and pairing type highlights opposing trends for SSB and non-SSB individuals. Lines represent predicted means from a Poisson GLMM fit by maximum likelihood (Laplace approximation), with shaded areas indicating 95% confidence intervals.

After being provided with a carcass, 5 of 83 male-female pairings (6%) were unsuccessful in laying eggs. Four out of 5 of these unsuccessful pairings were with an SSB individual (1 mounter, 3 mounted). In 24 of the 83 pairings (29%), the brood died before larval dispersal. SSB males were associated with 20 of these brood failures (12 mounter, 8 mounted). However, the difference in brood failure rate between SSB males and non-SSB males was not significant (Fisher’s exact test, *p* = 0.07).

### (b) Behaviour during male-male (MM) pairings

We modeled behavioural probability changes after the occurrence of SSB during male-male trials and contrasted this with temporal behavioural dynamics in non-SSB male-male trials using an anchor time of t = 10 min as described in the Methods (Equation 1; Figure 3A). Behavioural measures were grouped into categories representing aggressive behaviour (all instances of grappling or biting) and asocial behaviour (all instances of not-interacting or hiding). Both aggressive and asocial behaviour were reduced following same-sex mounting, but the reduction in asocial behaviour was more prominent. Asocial behaviour was also significantly higher in non-SSB individuals (mean = 20.5 counts) than in SSB individuals (mean counts = 10.0; Wilcoxon rank-sum test, p < 0.001; Figure 3C).

**Figure 3.**
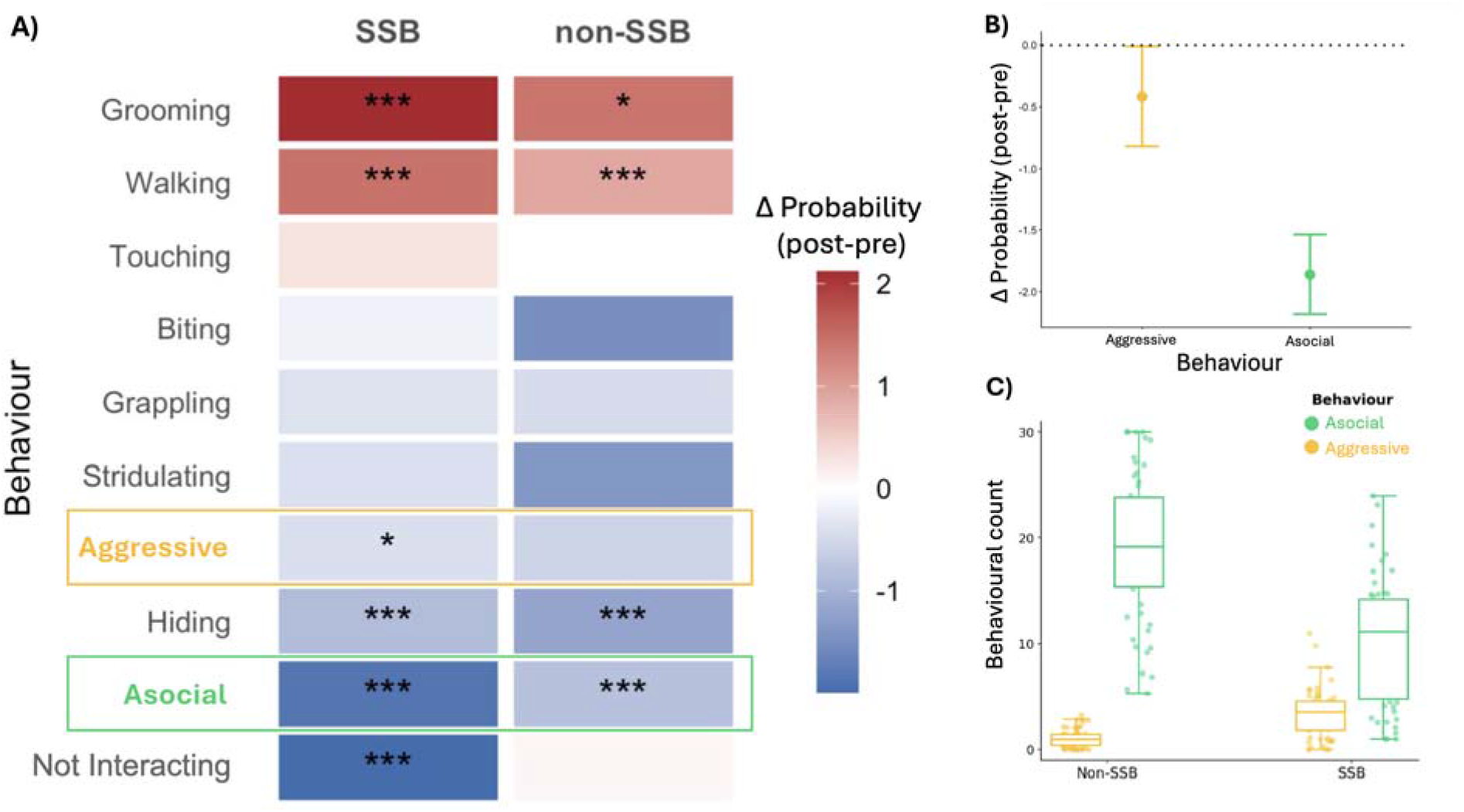
Behavioural changes after male-male mounting during male-male trials. (A) Heatmap showing the change in probability of behaviours expressed before and after male-male mounting. Columns represent whether the pair exhibited SSB or not. As no mounting occurred in non-SSB pairs, the non-SSB pre vs. post probabilities were calculated using an anchor time value as described in the Main Text. Higher Δ values mean that a behaviour is more likely to happen after mounting (or the anchor value) than before. Statistical significance is indicated for each Δ value: * = *p* < 0.05*, ** = p* < 0.01*, \*\*\** = *p* < 0.001. (B) Change in predicted probability of aggressive and asocial behaviour from pre-to post-mating for SSB individuals only. Points represent model-estimated Δ probabilities (post - pre) based on marginal means from GLMMs, and error bars indicate 95% CIs. The 95% CI on aggressive behaviour does not overlap 0. (C) Aggression and asocial behaviour by SSB status. Boxplot shows the distribution of behavioural counts for each group, with individual observations overlaid. Horizontal lines show medians, boxes show interquartile (IQR) ranges, and whiskers show extreme points within 1.5 x IQR.

A key finding was that not interacting was 88% less likely after mounting in SSB pairs (Δprob = −2.06, *p* < 0.001) but showed no difference before and after the anchor event for non-SSB pairs (Δprob = 0.08, *p* = 0.668). In general, asocial behaviour decreased after the mounting/anchor time for both SSB and non-SSB individuals, but the change in probability was significantly greater in SSB males than non-SSB males (z = −4.54, *p* < 0.001). In cases where asocial behaviour did occur in SSB males, it was significantly less likely to occur after mounting (*p* < 0.001; Figure 3B). Additionally, within SSB pairs the probability of displaying aggressive behaviour was reduced following mounting (Δprob = −0.42, *p* = 0.044), but there was no change around the anchor event for non-SSB pairs (Δprob = 0.56, *p* = 0.12; Figure 3B). Thus, same-sex mounting caused males to decrease the expression of aggressive behaviours.

The probability of walking and self-grooming was significantly increased following mounting (or anchor event) for both SSB males and non-SSB males, but the effect size was twice as large in SSB pairs for both behaviours. In SSB pairs, walking behaviour was 4.2 times more likely after mounting (Δprob = 1.45, *p* < 0.001) but only 2.1 times more likely after the anchor time for non-SSB pairs (Δprob=0.89, *p* < 0.001). Self-grooming was 8.4 times more likely after mounting in SSB pairs (Δprob= 2.12, *p* < 0.001) but only 4.3 times more likely after the anchor time in non-SSB pairs (Δprob = 1.43, *p* = 0.02). The difference in the change of probability after mounting/anchor between SSB and non-SSB individuals was significant for walking (z = 2.18, *p* = 0.03) but not for self-grooming (z = 0.82, *p* = 0.41). Thus, mounting prompted an increase in walking but not self-grooming behaviour.

### (c) Mounting behaviour during male-female (MF) pairings

SSB individuals mounted females less than non-SSB individuals overall (gaussian GLM, β = −5.47, t_79_ x= −2.53, *p* = 0.013). For non-SSB individuals, all males mounted the same amount regardless of their mass (β = −0.006, t_79_ = −0.74, p = 0.46), but for SSB males, heavier individuals mounted females more (β = 0.024, t_79_ = 2.40, *p* = 0.02). This pattern was especially prominent for males in the mounter role (β = 0.028, t_79_ = 2.38, p = 0.02), but less so for mounted males (β = 0.026, t_79_ = 1.98, *p* = 0.05). Male mass varied with role (one-way ANOVA, F_2, 80_ = 5.44, *p* = 0.006). Mounted individuals were significantly heavier than mounters (mean mass = 237.9 vs. 206.6 mg, respectively; Tukey HSD, *p* = 0.023), as well as non-SSB individuals (mean mass = 237.9 vs 202.3 mg, respectively; Tukey HSD, *p* < 0.01). However, male mass alone was not a significant predictor of mounting (Gaussian GLM, β = 0.01, t_79_ = 1.68, *p* = 0.097).

Males who exhibited SSB had similar mounting times in subsequent different-sex encounters compared with males who did not exhibit SSB (log-rank test, χ²(1) = 0.01, *p* = 0.94). Similarly, whether SSB males received or initiated SSB did not influence their mounting time with females (log-rank test, χ²(2) = 0.04, *p* = 0.98). The mass difference between males and females in each trial showed a marginally significant positive association with the likelihood of mounting for both SSB and non-SSB individuals (Cox proportional hazard, HR = 1.004, p=0.049). This means heavier individuals were more likely to mate with females at a faster speed than lighter individuals. However, the difference in mounting speed between SSB and non-SSB individuals, even when accounting for mass, was still nonsignificant (Cox proportional hazard; HR = 0.9995, p = 0.93).

### (d) Reproductive success

A composite measure of reproductive success was created using standardized z-scores from three variables with equal weighting: mean larval mass, brood size, and brood survival. The creation of this composite was justified by high internal consistency (Cronbach’s α = 0.89) and strong correlation between variables (average r = 0.73; Table S1). Two of the components, brood size and brood survival, were negatively correlated with SSB (Welch’s t_45.79_ = 2.89, *p* = 0.006; Welch’s t_51.4_ = 2.62, *p* = 0.012). The third component, mean larval mass, showed no correlation with SSB (Welch’s t_60.57_ = 1.03, *p* = 0.31).

Overall, non-SSB individuals showed higher reproductive success than SSB individual (Welch’s t_54.03_ = 2.51, *p* = 0.015; Figure 4A). Additionally, broods of non-SSB individual dispersed faster (Fine-Gray CRR; coef = −0.45, *p* = 0.046) (Figure 4C). There was no significant difference in dispersal time based on whether individuals were mounters, mounted, or did not express SSB (Fine-Gray CRR; coef = −0.55, *p* = 0.09), and prior SSB had no effect on other measures of fitness, such as the number of eggs laid (Welch’s t_57.86_ = 0.29, *p* = 0.77) or parental care duration (Welch’s t_50.31_ = 0.02, *p* = 0.98).

**Figure 4.**
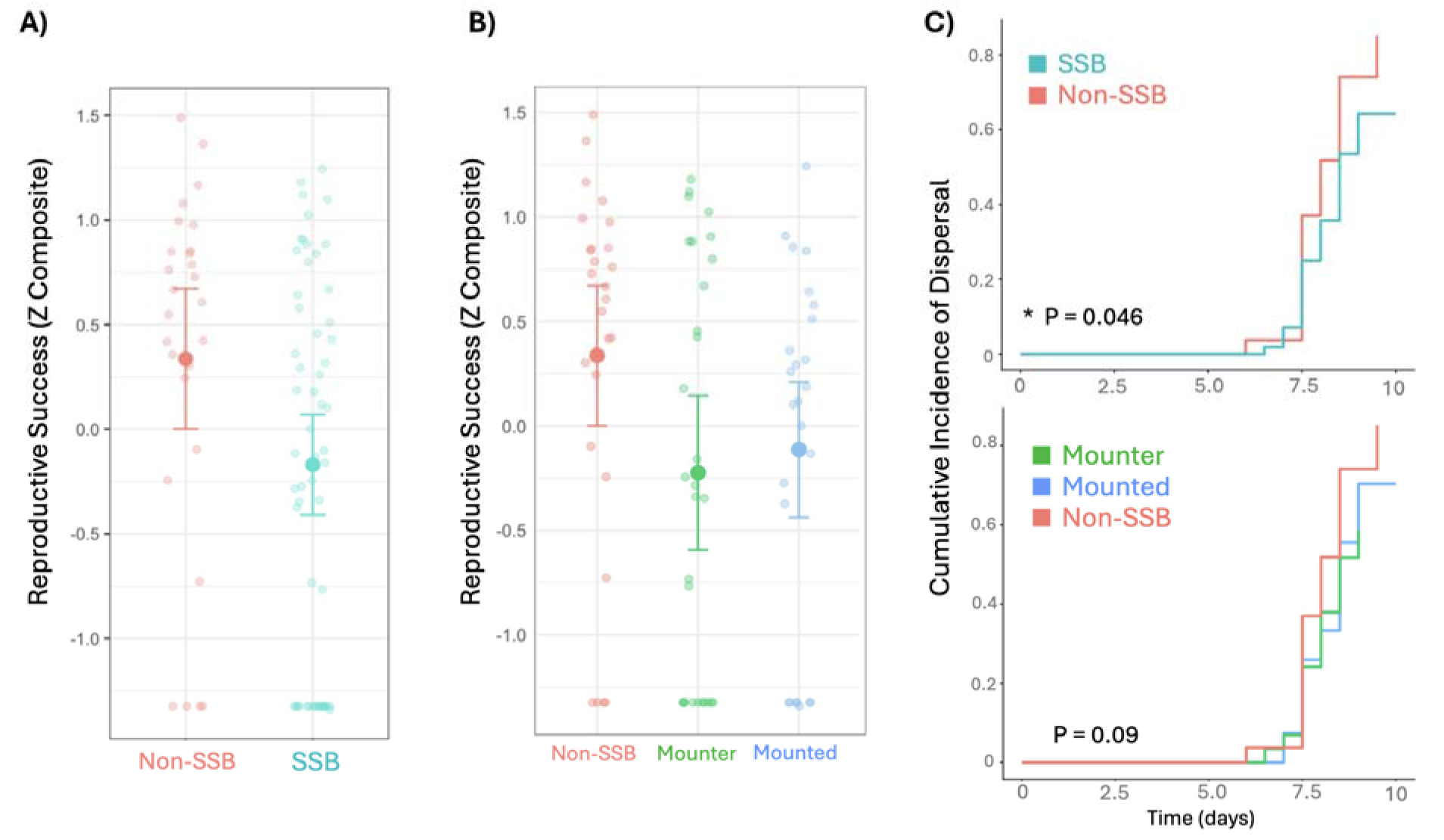
Fitness effects of SSB. (A) Reproductive success (z-scored composite) by SSB status. Median and interquartile range (B) Reproductive success by SSB role. Large solid circles indicate means and error bars represent 95% confidence intervals. (C) Cumulative incidence of dispersal over time by SSB status (above) and role (below), estimated using a Fine–Gray competing risks model (Competing event = death of brood).

## 4. Discussion

Taxonomic disparity in evolutionary explanations for SSB may reflect genuine taxonomic differences, but it must also be considered that systematic biases in our conceptions of complex animal behaviour could influence which hypotheses are tested and how the data are interpreted. Disentangling these requires attention to both adaptive and non-adaptive hypothesis testing in a broad range of animals. In invertebrates such as *N. vespilloides,* SSB has tended to be explained by non-adaptive hypotheses focused on the costs versus benefits of indiscriminate mating (e.g. Scharf & Martin, 2013; Engel *et al.,* 2015; Boutin *et al*., 2016), with studies often inferring that same-sex mating attempts represent mistakes due to inaccurate sex recognition (e.g. Sales *et al.,* 2018). However, more recent meta-analytical, comparative, and detailed neuroethological studies in insects have found that not only can SSB serve an adaptive function in the context of reducing aggression or providing partial fitness benefits when opposite-sex mates are scarce (Mizumoto *et al*., 2022; Green *et al.,* 2026) but also that SSB can be caused by a deliberate choice *despite* accurate sex recognition (Ouada *et al*., 2026).

Here, we found clear fitness costs associated with SSB, which does not concord with a common assumption of insect SSB research that this behaviour is a neutral, or low-cost outcome. In *N. vespilloides*, these costs are strongly driven by higher brood failure rates for males that had either initiated or been subjected to SSB in previous male-male encounters. However, we also found support for adaptive tension-reduction and dominance hypotheses; SSB appears to diminish costly aggressive behaviour during the course of ongoing male-male interactions. There was no evidence that it serves as practice to improve later different-sex mating encounters. The exact balance of these fitness costs and benefits is difficult to quantify, but the frequent occurrence of male SSB that reduces aggression, coupled with the clear sexually dimorphic pheromone profiles of these insects (Keppner *et al.,* 2017) suggest male SSB may be an evolutionary adaptation and not a simple byproduct of selection on indiscriminate mating.

We observed that more than 50% of male burying beetles engaged in SSB, which is consistent with previous studies in this and some other insects, such as the flour beetle *Tribolium castaneum* (Aiken, 1981; Serrano *et al.,* 1991, 2000; Engel *et al*., 2015; Morelle, *et al.,* 2026). Still, there are reasons to believe that this may be an underestimation of SSB in burying beetles. In male-male pairings, non-SSB individuals spent approximately two-thirds of the observation window during trials, either hiding or not interacting with the other individual. However, SSB individuals spent only a third of their trial, on average, not engaged in social behaviour, and in these individuals, almost all asocial behaviour occurred *before* mounting (Figure. 3). This indicates that once male beetles started interacting with each other, mounting was likely to follow, with a decreased likelihood of returning afterwards to asocial behaviour. In insects, same-sex mountings are normally briefer than different-sex matings (Benz, 1973; Henry, 1979). However, we found that mounting durations did not differ between same-sex and different-sex pairings, the proportion of males that mounted other males was the same as the proportion which mounted females, and rates of aggressive behaviour were the same for both male-female and male-male pairs.

Non-adaptive hypotheses hinge on the assumption that the cost of SSB is either non-existent or minor enough to not incur selection (Bailey and Zuk 2009; Lerch & Servedio, 2020). However, we found robust evidence that the expression of SSB lowers the reproductive success of burying beetles; this reduction in fitness was substantial, significantly decreasing brood size, brood survival, and larval dispersal probability (Figure 4). However, our findings also suggest that when male beetles do interact, aggression is likely, and in those cases where aggression occurs SSB can mitigate the conflict (Figure 3). In cases where SSB is not expressed, the high levels of asocial behaviour we found suggest that males reduce the cost of aggression by simply not engaging with the other male at all. Thus, SSB may represent one of a repertoire of behavioural choices that males can deploy to reduce the incidence of costly aggression. Similar trends have been observed in other insects. For example, in some field crickets (*Gryllus* spp.), SSB expressed by one individual in a pair was correlated with decreased aggression shown thereafter by the other individual (Green *et al.,* 2026). Here, we examined the impact of SSB on *both* individuals of the pair and still found a detectable tension-reduction effect: SSB was correlated with higher aggression, but our temporal analysis revealed that the first instance of mounting caused a significant decrease in the likelihood of aggressive behaviours thereafter (Figure 3B). Thus, SSB appears to manifest during, and subsequently reduce the intensity of, high-stakes aggressive encounters. While such a role for SSB during dynamic behavioural adjustment is relatively understudied in insects, it has been described in mammals, where SSB is associated with higher rates of aggression and conspecific killing (Goméz et al., 2023).

Roles (mounter vs. mounted) in all but one male-male pairing were fixed and unidirectional, though mounted individuals were heavier than both mounters and non-SSB individuals. This suggests that males seek individuals larger than them to mount, a pattern that is echoed in the literature (e.g. in the water bug *Palmacorixa nana*; Aiken, 1981) but is likely unrelated to sexual size dimorphism, as *N. vespilloides* is sexually monomorphic for size (Al Shareefi & Cotter, 2019; Sun et al., 2020). Lane *et al*. (2016) also found that 70% of male-male pairs had fixed roles in the flour beetle *Gnatocerus cornutus*, and they suggested that the purpose of these roles was to assert dominance. When *N. vespilloides* males fight over carcasses or access to female mating opportunities, larger individuals win fights more often (Otronen, 1988; Eggert et al., 2008; Lee et al., 2014). Thus, it is possible that smaller males view larger males as a potential threat, and similar to *G. cornutus,* attempt to win disputes by mounting and establishing dominance. The establishment of dominance hierarchies has a clear fitness benefit to male burying beetles, granting dominant males more access to females and increased paternity (House *et al.,* 2007; Müller *et al.,* 2007), and even in conditions where no carcass was present (such as this study), dominant males sire more offspring (Pettinger *et al.,* 2011).

Substantial fitness benefits such as we have found may provide insight into the evolutionary mechanisms by which SSB is maintained despite observed fitness costs. What ultimately matters is the net direct and indirect fitness associated with SSB expression; here, it seems plausible that SSB could be maintained because benefits of alleviating aggression outweight costs to reproductive output. In the future, studies that can integrate theses multifactorial costs and benefits across subjects’ entire lifespans would be helpful for evaluating this idea. It is increasingly understood that there is no one-size-fits-all explanation for SSB in animals. The trait, rather like other socio-sexual behaviours, may represent a complex evolutionary history with more than one explanation, even within a single lineage, and our findings support the value of testing multiple explanations for SSB and its fitness consequences in a combined framework.

## Ethics

Ethical approval was not required for this work.

## Data accessibility

All underlying data are available in the Supplementary Materials online.

## Declaration of AI use

Github copilot (an AI tool) was used to select suitable code for some statistical analyses (Binomial GLMM) and figure formatting (Figure 3). All code was manually checked.

## Authors’ contributions

N.E.P.: conceptualization, experimental design, data collection and curation, analysis, writing – original draft, writing – review and editing; J.W.: experimental design, data collection, writing – review and editing; N.P.: experimental design, funding acquisition, writing – review and editing; N.W.B.: funding acquisition, conceptualization, experimental design, writing – review and editing.

## Acknowledgements

We thank Rebekah Douglas, Solène Morelle, and Isobel Grieve for providing help with insect rearing and logistics in the laboratory. Thomas Green generously provided advice on earlier drafts of the manuscript.

