## Supplementary Material for "Same-sex sexual behaviour reduces reproductive success but mitigates aggression in a subsocial insect"

**Table S1:** Correlation matrix of the variables used in z-score composite measure of reproductive success

|  | Mean Larval Mass | Brood Size | Hatching Success |
| --- | --- | --- | --- |
| Mean Larval Mass | 1.000 | 0.638 | 0.700 |
| Brood Size | 0.638 | 1.000 | 0.841 |
| Hatching Success | 0.700 | 0.841 | 1.000 |

**Table S2:** PCA results showing standard deviation and proportion of variance explained for 3 proposed principal components (PC1, PC2, and PC3) used to evaluate reproductive success z-composite

|  | PC1 | PC2 | PC3 |
| --- | --- | --- | --- |
| Standard Deviation | 1.561 | 0.627 | 0.388 |
| Proportion of Variance | 0.818 | 0.132 | 0.051 |
| Cumulative Proportion | 0.818 | 0.949 | 1.000 |

**Table S3:** Loadings of each variable on the principal components described in Table S2

|  | PC1 | PC2 | PC3 |
| --- | --- | --- | --- |
| z_MeanLarvalMass | 0.540 | 0.828 | -0.151 |
| z_BroodSize | 0.586 | -0.499 | -0.639 |
| z_HatchingSuccess | 0.604 | -0.257 | 0.755 |

**Notes:** A principal components analysis (PCA) showed that all three reproductive traits loaded strongly on the first principal component, which explained 81.8% of the variance. The PCA-derived composite was strongly correlated with the z-score composite (r = 0.9998), confirming that using equal weighting of variables in the z-score composite was appropriate. Therefore, the z-score composite was used in subsequent analyses.

Raw behavioural, morphological, and reproductive success data are provided in the file “BB_SSB_Data.xls”. The file consists of 4 tabs: “RawBehaviouralData”, “MMPairings”, “MFPairings”, and “FormattedData”. The tab “RawBehaviourData” provides minute-by-minute data for every behaviour observed during both male-male and male-female pairings. Data are organised by focal individual and pair ID. The tab “MMPairings” provides summary data about the morphology of beetles in each pairing, the time and date of each pairing, as well as a summative count of all behaviours that occurred in each pairing. The tab “MFPairings” provides the same, but for Male-Female pairings, reproductive success data, and morphological data on individuals taken at the point when they were removed from their carcass (either due to death, dispersal, or removal after presence/absence observations were concluded). The tab “IndividualData” provides morphological, reproductive success, and behavioural count data for every individual.

Tab1: “RawBehaviourData”

| **Heading** | **Title** | **Description** |
| --- | --- | --- |
| 1 | Pair_ID | Unique number assigned to each pairing  (“weekofexperiment”.”dayofexperiment”.”numberofpairing”) |
| 2 | Minute | The minute of each 30-minute pairing in which the observation occurred |
| 3 | Individual_ID | Unique number assigned to each test subject |
| 4 | Role | Individual identifier for male-male pairings  For male-male pairings:  (1 = the male with Tipp-Ex applied, 2 = the male with no Tipp-Ex applied)  For male-female pairings:  (1 = male, 2 = female) |
| 5 | MM | Whether the pairing was male-male or not  (0 = negative, 1 = affirmative) |
| 6 | MF | Whether the pairing was male-female or not  (0 = negative, 1 = affirmative) |
| 7 | Grappling | Whether the individual in that pair displayed grappling behaviour during that minute  (0 = negative, 1=affirmative) |
| 8 | Mounting | Whether the individual in that pair displayed mounting behaviour during that minute  (0 = negative, 1=affirmative) |
| 9 | Touching | Whether the individual in that pair displayed touching behaviour during that minute  (0 = negative, 1=affirmative) |
| 10 | Walking | Whether the individual in that pair displayed walking behaviour during that minute  (0 = negative, 1=affirmative) |
| 11 | Biting | Whether the individual in that pair displayed biting behaviour during that minute  (0 = negative, 1=affirmative) |
| 12 | Stridulating | Whether the individual in that pair displayed stridulating behaviour during that minute  (0 = negative, 1=affirmative) |
| 13 | Not_interacting | Whether the individual in that pair displayed not-interacting behaviour during that minute  (0 = negative, 1=affirmative) |
| 14 | Selfgrooming | Whether the individual in that pair displayed wing movement or self-grooming behaviour during that minute  (0 = negative, 1=affirmative) |
| 15 | Hiding | Whether the individual in that pair displayed hiding behaviour during that minute  (0 = negative, 1=affirmative) |
| 16 | Mating_attempt | Whether the individual in that pair displayed a failed mating attempt during that minute  (0 = negative, 1=affirmative) |
| 17 | Behaviour | The unique code associated with the behaviour displayed during that minute  g = grappling  m = mounting  t = touching with antennae  w = walking  b = biting  s = stridulating  n = not-interacting  e = wing movement or self-grooming  h = hiding |
| 18 | Behaviour2 | If two behaviours were observed in a minute, the second behaviour was listed here |
| 19 | Behaviour3 | If three behaviours were observed in a minute, the third behaviour was listed here |
| 20 | StartOfMountingBout | A marker to denote whether that minute was the first minute in a multi-minute mounting bout (i.e. to separate out discrete mounting occurrences)  (0 = negative, 1 = affirmative) |

Tab 2: “MMPairings”

| **Heading** | **Title** | **Description** |
| --- | --- | --- |
| 1 | PairID | Unique number assigned to each pairing  (“weekofexperiment”.”dayofexperiment”.”numberofpairing”) |
| 2 | Male1ID | The unique identification number associated with male 1 of the pairing. Male 1 is the individual with Tipp-Ex applied |
| 3 | Male2ID | The unique identification number associated with male 2 of the pairing. Male 2 is the individual without Tipp-Ex applied |
| 4 | Male1Age | The age of male 1  (days since eclosion) |
| 5 | Male2Age | The age of male 2  (days since eclosion) |
| 6 | AgeDifference | The difference in age between the two males  (Male1Age - Male2Age) |
| 7 | AbsAgeDiff | The absolute value of the age difference between the two males |
| 8 | DatePaired | The date on which the pairing occurred |
| 9 | TimePaired | The time at which the pairing occurred |
| 10 | Male1Pronotum | A measure of the pronotum width of male 1  (mm) |
| 11 | Male2Pronotum | A measure of the pronotum width of male 2  (mm) |
| 12 | Male1Weight | The mass of male 1  (mg) |
| 13 | Male2Weight | The mass of male 2  (mg) |
| 14 | WeightDiff | The difference in mass between male 1 and male 2 (Male1Weight - Male2Weight) |
| 15 | PronotumDiff | The difference in pronotum width between male 1 and male 2  (Male1Pronotum - Male2Pronotum) |
| 16 | SSB | Whether the individual exhibited SSB  (0 = non-SSB, 1 = SSB) |
| 17 | Male1Mount | Whether male 1 mounted male 2  (0 = negative, 1 = affirmative) |
| 18 | Male2Mount | Whether male 2 mounted male 1  (0 = negative, 1 = affirmative) |
| 19 | Grappling | The sum of grappling behaviours displayed by both male 1 and male 2 during the 30-minute pairing |
| 20 | Mounting | The sum of mounting behaviours displayed by both male 1 and male 2 during the 30-minute pairing |
| 21 | Touching | The sum of touching behaviours displayed by both male 1 and male 2 during the 30-minute pairing |
| 22 | Walking | The sum of walking behaviours displayed by both male 1 and male 2 during the 30-minute pairing |
| 23 | Biting | The sum of biting behaviours displayed by both male 1 and male 2 during the 30-minute pairing |
| 24 | Stridulating | The sum of stridulating behaviours displayed by both male 1 and male 2 during the 30-minute pairing |
| 25 | Not_Interacting | The sum of not-interacting behaviours displayed by both male 1 and male 2 during the 30-minute pairing |
| 26 | Self_Grooming | The sum of wing movement or self-grooming behaviours displayed by both male 1 and male 2 during the 30-minute pairing |
| 27 | Hiding | The sum of hiding behaviours displayed by both male 1 and male 2 during the 30-minute pairing |
| 28 | Mating_Attempt | The sum of failed mating-attempts displayed by both male 1 and male 2 during the 30-minute pairing |

Tab3: “MFPairings”

| **Heading** | **Title** | **Description** |
| --- | --- | --- |
| 1 | PairID | Unique number assigned to each pairing  (“weekofexperiment”.”dayofexperiment”.”numberofpairing”) |
| 2 | MaleID | The unique identification number associated with the male of the pairing |
| 3 | FemaleID | The unique identification number associated with the female of the pairing |
| 4 | MaleEclosionDate | The date on which the male eclosed |
| 5 | FemaleEclosionDate | The date on which the female eclosed |
| 6 | MaleAge | The age of the male  (days since eclosion) |
| 7 | FemaleAge | The age of the female  (days since eclosion) |
| 8 | AgeDifference | The difference in age between the two individuals  (MaleAge- FemaleAge) |
| 9 | AbsAgeDiff | The absolute value of the age difference between the male and female |
| 10 | DatePaired | The date on which the pairing occurred |
| 11 | TimePaired | The time at which the pairing occurred |
| 12 | DateRemoved | The date on which the male was removed from the phase 3 box, either in the morning (AM) or the evening (PM). |
| 13 | FirstAbsence | The date on which the male’s first absence from the crypt was recorded during presence/absence observations.  “dispersed” = male was present until larvae dispersed  “DEAD” = male died  “Infanticide” = male was removed immediately due to infanticide and thus had no first absence  “No eggs laid” = the pairing did not lay any eggs  “Unhatched” = eggs were laid but they did not hatch  “Date + Infanticide” = the male was absent from the crypt at least once prior, but then was removed early due to infanticide |
| 14 | MalePronotum | The pronotum width of the male  (mm) |
| 15 | MalePreWeight | The mass of the male prior to male-male pairing  (mg) |
| 16 | FemalePronotum | The pronotum width of the female  (mm) |
| 17 | FemaleWeight | The mass of the female prior to male-female pairing  (mg) |
| 18 | MaleFinalWeight | The mass of the male after the experiment was finished and all larvae reached dispersal, or whenever the male was removed early due to premature death, infanticide, or prolonged absence from the crypt.  (mg) |
| 19 | MaleWeightChange | The overall change in mass between the beginning of the experiment and the end of the experiment  (MaleFinalWeight - MalePreWeight) |
| 20 | PartnerWeightDiff | The difference in mass between the and the female (MalePreWeight - FemaleWeight) |
| 21 | PronotumDiff | The difference in pronotum width between the male and the female  (MalePronotum - FemalePronotum) |
| 22 | DispersalDate | The date on which the larvae dispersed, either in the morning (AM), or the evening (PM). |
| 23 | MaleSurvivalToDispersal | Whether the male survived until its larvae dispersed  (0 = negative, 1 = affirmative) |
| 24 | SSB | Whether the male had previously exhibited SSB in male-male pairings  (0 = negative, 1 = affirmative) |
| 25 | Mounter | Whether the male had mounted another male in male-male pairings  (0 = negative, 1 = affirmative) |
| 26 | Mounted | Whether the male had been mounted by another male in male-male pairings  (0 = negative, 1 = affirmative) |
| 21 | Grappling | The sum of grappling behaviours displayed by both the male and female during the 30-minute pairing |
| 22 | Mounting | The sum of mounting behaviours displayed by both the male and female during the 30-minute pairing |
| 23 | Touching | The sum of touching behaviours displayed by both the male and female during the 30-minute pairing |
| 24 | Walking | The sum of walking behaviours displayed by both the male and female during the 30-minute pairing |
| 25 | Biting | The sum of biting behaviours displayed by both the male and female during the 30-minute pairing |
| 26 | Stridulating | The sum of stridulating behaviours displayed by both the male and female during the 30-minute pairing |
| 27 | Not_Interacting | The sum of not-interacting behaviours displayed by both the male and female during the 30-minute pairing |
| 28 | Self_Grooming | The sum of wing movement or self-grooming behaviours displayed by both the male and female during the 30-minute pairing |
| 29 | Hiding | The sum of hiding behaviours displayed by both the male and female during the 30-minute pairing |
| 30 | Mating_Attempt | The sum of failed mating-attempts displayed by both the male and female during the 30-minute pairing |

Tab 4: “IndividualData”

| **Heading** | **Title** | **Description** |
| --- | --- | --- |
| 1 | Individual_ID | Unique number assigned to each test subject |
| 2 | PairType | Whether the trial was male-male (MM) or male-female (MF) |
| 3 | Role | Individual identifier for male-male pairings  (Male1 = the male with Tipp-Ex applied, Male2 = the male with no Tipp-Ex applied) |
| 4 | Pronotum | A measure of the pronotum width of each individual (mm) |
| 5 | MMPronotumDiff | The difference in pronotum width between the males in each male-male trial.  (Male 1 Pronotum Width - Male 2 Pronotum Width) |
| 6 | Mass | The mass of the individual.  Males: measured immediately prior to the male-male pairing  Females: measured immediately prior to the male-female pairing |
| 7 | MMWeightDiff | The difference in mass between the males in each male-male trial.  (Male 1 Mass - Male 2 Mass) |
| 8 | MFWeightDiff | The difference in mass between the male female in each male-female trial  (Male Mass - Female Mass) |
| 9 | Age | The age of each individual at the time of first pairing  (Number of days since eclosion) |
| 10 | WeightChange | The difference in the mass of an individual over the course of the experiment. The first measurement was taken immediately prior to male-male pairings. The second measurement was taken when the individual was removed from their reproductive box at the end of phase 3 of the experiment.  (First measurement - Second measurement) |
| 11 | SSB | Whether the individual exhibited SSB  (0 = non-SSB, 1 = SSB) |
| 12 | Mounter | Whether the individual was a “mounter” or not. I.e. whether the individual mounted the other male in a male-male pairing where SSB occurred  (0 = non-mounter, 1 = mounter) |
| 13 | Mounted | Whether the individual was a “mounted” or not. I.e. whether the individual was mounted by the other male in a male-male pairing where SSB occurred  (0 = non-mounter, 1 = mounter) |
| 14 | MMTimeToMounting | The minute at which the first instance of mounting was observed in male-male pairings |
| 15 | MMSumMounting | The total number of minutes in which mounting was displayed by the individual in male-male pairings |
| 16 | MMSumMountingBouts | The number of unique, discrete mounting events displayed by the individual in male-male pairings, referred to as bouts  (Instances of mounting separated by at least one minute of a different behaviour) |
| 17 | MMAvMountBoutDuration | The average duration of each bout of mounting in male-male pairings  (Minutes) |
| 18 | MFTimeToMounting | The minute at which the first instance of mounting was observed in male-female pairings |
| 19 | MFSumMounting | The total number of minutes in which mounting was displayed by the individual in male-female pairings |
| 20 | MFSumMountingBouts | The number of unique, discrete mounting events displayed by the individual in male-female pairings, referred to as bouts  (Instances of mounting separated by at least one minute of a different behaviour) |
| 21 | MFAvMountBoutDuration | The average duration of each bout of mounting in male-female pairings  (Minutes) |
| 22 | NumEggsLaid | The amount of eggs visible on the underside or walls of the phase 3 box, counted 2 days after the pairing had been given a carcass. |
| 23 | BroodSize | The number of larvae which survived to dispersal |
| 24 | HatchingSuccess | The proportion of eggs counted which successfully hatched and survived to dispersal  (BroodSize / NumEggsLaid) |
| 25 | Infanticide | Whether the individual displayed infanticidal behaviour prior to removal  (0 = no infanticidal behaviour, 1 = infanticidal behaviour) |
| 26 | ParentalCareDuration | The length of time that the individual was displaying parental care behaviour to its larvae. Measured from 2 days after carcass was provided in phase 3, to the day of first absence)  (days) |
| 27 | BroodMass | The total mass of all the larvae which survived to dispersal  (mg) |
| 28 | MeanLarvalMass | The average mass of individual larva  (BroodMass / BroodSize)  (mg) |
| 29 | TimeToDispersal | The length of time that the larvae took to reach dispersal after presence/absence observations had begun  (days) |
| 30 | CareToDispersal | The proportion of the larvae’s developmental period that the individual was providing parental care  (ParentalCareDuration / TimeToDispersal) |
| 31 | LarvalSurvivalToDispersal | Whether the larvae survived to dispersal or not  (0 = none survived, 1 = one or more larva survived) |
| 32 | CarcassWeight | The mass of the mouse carcass provided to the individual during phase 3  (g) |
